# Developmental Programming of Mitochondrial Function Limits Lifespan in Short-Lived Animals

**DOI:** 10.1101/2023.06.23.546283

**Authors:** Beatriz Castejon-Vega, Ignacio Fernandez-Guerrero, Yizhou Yu, Rachel Zussman, Tetsushi Kataura, Rhoda Stefanatos, Sarah Allison, Davide D’Andrea, Mario Cordero, Milos Filipovic, Miguel Martins, Viktor I. Korolchuk, Alberto Sanz

## Abstract

The contribution of mitochondria to lifespan determination remains controversial, as impaired mitochondrial function can paradoxically both shorten and extend longevity. During ageing, mitochondria accumulate defects that disrupt electron transport and elevate the production of reactive oxygen species (ROS) per unit of ATP. Here, we developed *Drosophila melanogaster* models carrying mitochondria that phenocopy aged organelles—termed “aged-like” mitochondria—to dissect the developmental versus adult contributions of mitochondrial dysfunction to lifespan regulation. Inducing aged-like mitochondria during development caused profound metabolic maladaptation and markedly reduced adult lifespan, without signs of accelerated ageing. In contrast, restricting their expression to adulthood resulted in only a modest reduction in lifespan, accompanied by an increased mortality rate, indicative of accelerated ageing. Enhancing mitochondrial function exclusively during development by expressing the alternative oxidase (AOX) mitigated these metabolic defects and significantly extended adult survival. Likewise, developmental overexpression of Rheb, an activator of Target of Rapamycin (TOR) signalling, improved adult survival without restoring mitochondrial respiration. Finally, we show that mitochondrial respiratory capacity cannot be reinstated in adults with “aged-like” mitochondria, as oxidative phosphorylation (OXPHOS) protein levels are largely established during development and remain stable throughout adult life. We propose that *Drosophila* permanently tunes adult metabolism according to developmental cues to optimise reproductive fitness, at the expense of long-term survival.

## 1. Introduction

Mitochondria play a central role in determining cellular fate and are thought to exert a significant influence on organismal lifespan, although the precise nature of this relationship remains poorly defined (Lopez-Otin, Blasco et al. 2023). Dysfunctional mitochondria accumulate with age and are commonly observed in the cells of individuals with degenerative diseases such as Parkinson’s, Alzheimer’s, and cancer (Wallace 2005). These mitochondria exhibit reduced respiratory efficiency and generate increased levels of ROS (Graham, Stefanatos et al. 2022, Rimal, Tantray et al. 2023). Likewise, mutations in genes encoding mitochondrial proteins can cause severe mitochondrial disorders that markedly reduce lifespan (Gorman, Schaefer et al. 2015). Paradoxically, inhibition of mitochondrial function has been shown to mitigate the harmful effects of cellular senescence in mammalian cells and extend lifespan in model organisms ranging from *Caenorhabditis elegans* to mice (Dillin, Hsu et al. 2002, Dell’agnello, Leo et al. 2007, Copeland, Cho et al. 2009, Correia-Melo, Marques et al. 2016).

To reconcile these seemingly contradictory findings, we generated *Drosophila melanogaster* models in which mitochondrial function is selectively disrupted during development and adulthood, or exclusively during adulthood. We initially focused on mitochondrial complex I (CI) and demonstrated that developmental depletion of CI results in short-lived flies with severe metabolic dysfunction (Stefanatos, Robertson et al. 2025). In contrast, restricting CI depletion to adulthood produces long-lived flies without notable molecular or physiological abnormalities. Although both short- and long-lived CI-depleted flies replicate and help explain the opposing results reported in the literature— namely, lifespan shortening (Foriel, Renkema et al. 2019) and extension following CI depletion (Owusu-Ansah, Song et al. 2013) —they do not represent a suitable model for aged mitochondria. We and others have shown that depletion of CI disrupts oxidative phosphorylation (OXPHOS) without increasing ROS levels (Graham, Stefanatos et al. 2022, Granat, Ranson et al. 2022, Rimal, Tantray et al. 2023). However, aged mitochondria are characterised by high levels of mtROS (Cocheme, Quin et al. 2011, Scialo, Sriram et al. 2016, Graham, Stefanatos et al. 2022), a phenotype observed when CIV is depleted, but not when CI is depleted (Graham, Stefanatos et al. 2022).

Here, we disrupt complex IV (CIV) using RNA interference (RNAi) to generate ’aged-like’ mitochondria that mimic the mitochondrial phenotype observed in elderly individuals, characterised by reduced ATP production and increased generation of ROS per unit of ATP synthesised (Graham, Stefanatos et al. 2022). As expected, we observe both shared and distinct responses to CI and CIV depletion. While CI is required for lifespan regulation only during development, CIV influences lifespan through two distinct "windows of opportunity": the first, shared with CI, occurs during development, and the second, unique to CIV, emerges during senescence.

We demonstrate the importance of the developmental "window of opportunity" by rescuing adult lifespan through two distinct interventions applied exclusively during development. First, we complement CIV function using an alternative oxidase (AOX), which mitigates many of the metabolic maladaptations observed in CIV-depleted flies and extends lifespan. Second, we overexpress Rheb, an activator of the Target of Rapamycin (Tor) pathway, which prolongs lifespan independently of changes in mitochondrial respiration. These findings support the idea that mitochondrial function is essential for the proper adult metabolic programming that occurs during development.

Finally, we show that restoring mitochondrial activity in adults lacking CIV during development is not feasible, as the levels of CIV and other OXPHOS components are established during development and remain fixed throughout the fly’s reproductive lifespan. We propose that this developmental "fixing" of OXPHOS levels in response to environmental cues, along with the subsequent metabolic rewiring, represents an adaptation in short-lived postmitotic organisms such as *Drosophila melanogaster*. This strategy prioritises reproductive success at the expense of adult survival.

## 2. Materials and Methods

### Analysis of transcriptomic and proteomic data from FlyBase

Transcriptomic and proteomic expression data across *Drosophila melanogaster* developmental stages were retrieved from FlyBase (https://flybase.org). For mRNA expression, the *modENCODE mRNA-Seq Development* dataset (Brown, Boley et al. 2014) was used, comprising normalised RPKM values spanning embryonic to adult stages. Protein abundance was obtained from the *Development Proteome (Casas-Vila Proteome Life Cycle)* dataset (Casas-Vila, Bluhm et al. 2017), which provides label-free mass spectrometry-based quantification across matched developmental stages. For both datasets, developmental stages were grouped into 15 matched time points from 0–2 h embryos to 5-day-old adults. Biological replicates were averaged, and gene expression values were z-score normalised across developmental time for each gene independently.

Data processing and visualisation were performed in R using the tidyverse, svglite, and ggplot2 packages. Each dataset was reshaped into long format and tagged by data type (Transcript or Proteomic). For selected functional gene groups (i.e., OXPHOS subunits excluding testis-specific isoforms), both individual gene expression profiles and group-level averages were plotted. Expression trends were visualised as line plots with colour-coded traces for transcriptomic and proteomic data. Background gene-level trajectories were plotted with low opacity, while group mean trajectories were overlaid as bold dashed lines.

### Drosophila Activity Monitor (DAM) Adapted for Starvation Survival Stress

Starvation survival activity was also measured using the Drosophila Activity Monitor (DAM; Trikinetics, Waltham, MA, USA) equipped with 15 infrared beams per tube for precise movement detection. For each experimental condition, 32 individual flies were placed in separate 5 mm-diameter glass tubes containing 1% agar at one end (to provide moisture) and sealed with a cotton plug at the other.

Each monitoring unit was connected to a computer that continuously recorded activity, registering a movement event each time a fly crossed an infrared beam. Data were collected at 1-minute intervals, generating a detailed temporal record of locomotor activity and survival.

The units were maintained in a 25 °C incubator under constant environmental conditions and monitored until all flies had died. Time of death for each fly was defined as the last time point at which movement was detected.

### Fly Husbandry

*Drosophila melanogaster* flies were reared on a standard medium containing 1% agar, 1.5% sucrose, 3% glucose, 3.5% dried yeast, 1.5% maize, 1% wheat germ, 1% soybean flour, 3% treacle, 0.5% propionic acid, and 0.1% Nipagin. Unless stated otherwise, flies were maintained at 25 °C under a 12-hour light/dark cycle. Male flies were used for all experiments. Experimental cohorts were collected within 48 hours post-eclosion using CO₂ or ice anaesthesia and maintained at a density of 20 flies per vial.

To drive the expression of RNAi and transgenes, either the GeneSwitch (GS) system was employed in combination with varying concentrations of RU-486, or the GAL4/UAS system was used in conjunction with a temperature-sensitive GAL80 (tsGAL80) to allow temporal control of gene expression at specific developmental stages, such as the pupal stage when flies do not feed. The genotypes and sources of all fly strains used are listed in Supplementary Table 1.

### Generation of Transgenic Flies

To generate an RNAi-resistant transgene, we introduced silent mutations into the coding sequence of the COX5B cDNA. The modified cDNA was synthesised and subcloned by VectorBuilder into the pUASTattB-5xUAS/mini-Hsp70 transformation vector. UAS-Rheb was created in the same way using the non-mutated cDNA. Site-specific integration into the fly genome was performed by BestGene Inc. using the ΦC31 integrase system, targeting a defined intergenic landing site characterised by moderate transcriptional activity. Transformants were identified by the presence of the mini-white marker (eye colour), and homozygous stocks were established through standard genetic crosses. The resulting transgenic flies were crossed with those carrying a UAS-RNAi construct targeting COX5B to generate a stable fly stock harbouring both the RNAi and the RNAi-resistant rescue transgene.

### High-Resolution Respirometry

Mitochondrial respiration was assessed using an Oxygraph-2k (Oroboros Instruments) as previously described (Stefanatos, Robertson et al. 2025). Ten ∼5-day-old adult flies were homogenised in isolation buffer containing 250 mM sucrose, 5 mM Tris-HCl, and 2 mM EGTA (pH 7.4), and the homogenate was subsequently diluted 10-fold in assay buffer consisting of 120 mM KCl, 5 mM KH₂PO₄, 3 mM HEPES, 1 mM EGTA, 1 mM MgCl₂, and 0.2% (w/v) BSA (pH 7.2).

To assess CI-linked respiration, 5 mM proline and 5 mM pyruvate were added to the chamber, and state 3 respiration was initiated by the addition of 1 mM ADP. Complex IV (CIV)-linked respiration was measured using 0.5 mM N,N,N′,N′-tetramethyl-p-phenylenediamine (TMPD) and 2 mM ascorbate as electron donors, and 50 mM sodium azide was used to inhibit CIV.

Oxygen consumption rates were recorded using DatLab 7.0 (Oroboros Instruments) and normalised to the protein content of each homogenate, expressed as picomoles of O₂ per second per milligram of protein. Protein concentrations were determined using the Bradford assay.

### Lipid Staining and Confocal Microscopy

Fat bodies from ∼5-day-old flies were dissected and fixed in 4% paraformaldehyde for 30 minutes at room temperature. Following fixation, tissues were rinsed in 1× PBS and incubated for 1 hour at room temperature in a staining solution consisting of 1× PBS with 0.005% saponin, LipidTOX™ Red (1:500 dilution), and DAPI (1:1000 dilution). Samples were then washed three times for 15 minutes each in 1× PBS containing 0.005% saponin.

Tissues were mounted on polylysine-coated glass slides using 13 mm × 0.12 mm spacers and ProLong™ Diamond Antifade Mountant. Samples were imaged using a Leica SP8 DLS confocal microscope, and images were processed using ImageJ software.

### Proteomics

Proteomic analysis was performed at the MRC Toxicology Unit, University of Cambridge, as described in (Stefanatos, Robertson et al. 2025) .Briefly, 50 five-day-old male flies were homogenised in 100 mM TEAB buffer, supplemented with 1% RapiGest, and centrifuged. Proteins were reduced, alkylated, and digested with trypsin. Peptides were quantified and labelled using 2xTMT11 with pooled reference standards. LC-MS/MS was conducted on an Orbitrap Eclipse™ mass spectrometer using FAIMS and SPS-MS3 with real-time search. Data were analysed in Proteome Discoverer v2.5 using Sequest HT against the *Drosophila* Uniprot database, including common contaminants. Differentially expressed proteins were identified and subjected to functional enrichment analysis using the STRING database (Szklarczyk, Kirsch et al. 2023).

### Protein Turnover in Adult Flies

Protein turnover data for *Drosophila melanogaster* brain and thorax tissues were obtained from previously published datasets (Vincow, Thomas et al. 2021, Ren, Xu et al. 2023). To categorise proteins by subcellular localisation, orthologue-based annotations derived from human protein data available at the Human Protein Atlas (https://www.proteinatlas.org/about/download) were used. Briefly, human–fly orthologues were identified using gProfiler, and human protein localisation data were retrieved from the Human Protein Atlas. These annotations were subsequently merged with *Drosophila* gene identifiers to assign subcellular compartments to fly proteins. Data integration and analysis were performed in R. Additionally, oxidative phosphorylation (OXPHOS) subunit annotations were incorporated to determine whether their turnover rates differed significantly from other mitochondrial proteins. Protein half-life values were log-transformed, and differences across subcellular compartments as well as OXPHOS versus non-OXPHOS proteins were visualised using boxplots.

### qPCR gene expression analysis

Total RNA was extracted from ∼5-day-old adult flies using a standard Trizol-based method as previously described (Stefanatos, Robertson et al. 2025). RNA samples were treated with DNase I and purified by ethanol precipitation. Complementary DNA (cDNA) was synthesised using the High-Capacity cDNA Reverse Transcription Kit following the manufacturer’s instructions. Quantitative PCR was performed using QuantiNova SYBR Green reagents on an Applied Biosystems QuantStudio 3 instrument. Data were acquired and analysed using the QuantStudio Design & Analysis software. Relative expression levels of target genes were normalised to appropriate housekeeping genes (see Supplementary Table 1 for primer details).

### RNA Sequencing and Differential Expression Analysis

Total RNA was extracted as previously described for qPCR gene expression analysis and submitted to the Department of Biochemistry, University of Cambridge, for quality control, poly(A) selection, and library preparation using the TruSeq Stranded mRNA Library Prep kit. Sequencing was performed on the Illumina NextSeq 2000 platform using the NextSeq 2000 100-cycle (2×50 bp) P3 kit. Read quality was assessed with FastQC v0.11.9, and Kallisto v0.48.0 was used to create an alignment index based on the *Drosophila melanogaster* genome (BDGP6.32) and to map reads to the reference transcriptome. Data were imported into R using the rhdf5 v2.44.0 and tidyverse v2.0.0 packages, with Kallisto abundance files imported via tximport v1.28.0 and transcript annotations obtained using biomaRt v2.56.0. Low-count transcripts were removed, and normalisation was performed using edgeR v3.42.2. Principal component analysis (PCA) was conducted to identify the components explaining the greatest variance. Differential gene expression (DGE) analysis was performed using Limma v3.56.1, with DEGs selected based on a fold change (FC) threshold of ≥1.5 or ≤–1.5 and a false discovery rate (FDR) <0.05 unless otherwise stated. Functional enrichment analysis was carried out using the GSEABase v16.0, clusterProfiler v4.8.1, and gprofiler2 v0.2.1 packages.

### Survival Analysis

*Lifespan at 25°C*: Flies were transferred to fresh food every 2–3 days, and mortality was recorded at each transfer. *Hydrogen Peroxide Treatment*: Flies aged 3–5 days were placed in vials containing 5 ml of 1.5% agar, 5% sucrose, and 2.5% H₂O₂. Survival was monitored at regular intervals. *Thermal Stress:* Flies were maintained at 32°C and transferred to fresh media every 2–3 days, with mortality assessed throughout. *Starvation Stress:* Flies aged 3–5 days were placed in vials containing 2 ml of 1% agar. Deaths were recorded at consistent time points. Each experimental group consisted of a minimum of 100 flies, with experiments independently repeated 2–3 times unless otherwise stated.

### Statistical Analysis

Data visualisation was performed using GraphPad Prism 10 or R, depending on the specific analytical requirements. Statistical analyses included Analysis of Variance (ANOVA) or t-tests, chosen based on data distribution and experimental design. Post-hoc analyses were conducted following significant ANOVA results to identify differences between specific groups.

Survival curves were generated using GraphPad Prism 10 and analysed with the log-rank (Mantel–Cox) test. Mortality rates (µ(x)) were modelled using the Gompertz equation (µ(x)=αe^βx^) by fitting age-specific death data with a robust maximum-likelihood approach. For each experimental group, the Gompertz parameters α (baseline mortality rate) and β (rate of age-related mortality increase) were estimated using bounded optimisation with multiple initial values to ensure convergence, and 95% confidence intervals were derived from numerical differentiation.

Statistical significance was defined as *p* < 0.05, and data are presented as mean ± SEM unless stated otherwise.

### Triacylglycerol (TAG) measurement

For each sample, 15 adult male flies were weighed and homogenised in Tet buffer (10 mM Tris, 1 mM EDTA, 0.1% v/v Triton-X-100, pH 8.0) using glass beads and an OMNI Bead Ruptor. Homogenates were incubated at 72°C for 20 min, followed by centrifugation at 13,000 × g for 1 min at 4°C. Supernatants were further incubated at 72°C for 15 min. Subsequently, 3 µl aliquots of samples or glycerol standards were mixed with 300 µl ThermoFisher Infinity Triglyceride reagent and incubated for 15 min at 37°C. Absorbance was measured at 540 nm using a Multiskan FC plate reader (ThermoFisher). TAG concentrations were calculated from the glycerol standard curve and normalised by fly weight, expressed as µg TAG per mg fly.

### Western blot

Three to four replicates of 10 flies per condition were lysed in RIPA buffer (Sigma-Aldrich) supplemented with Halt Protease and Phosphatase Inhibitor Cocktail (ThermoFisher Scientific), using a motorised pestle. Lysates were centrifuged at 13,000 × g for 10 min at 4°C, and protein concentrations in the supernatants were measured using the DC Protein Assay (BioRad) and a GloMax plate reader (Promega). Samples were prepared by boiling at 100°C for 5 min in 2× Laemmli sample buffer (BioRad) containing 2.5% β-mercaptoethanol (Sigma-Aldrich). Equal amounts of protein (40 µg) were separated by SDS-PAGE using 8% or 15% Tris-glycine gels and transferred onto Immobilon-P PVDF membranes (Millipore) with a TransBlot SD Semi-Dry Transfer Cell (BioRad). Membranes were blocked with 5% milk in PBS-Tween (PBST) for 30 min at room temperature, then incubated overnight at 4°C with primary antibodies diluted in 5% BSA or 5% milk containing 0.02% sodium azide. Secondary HRP-conjugated antibodies (anti-rabbit or anti-mouse; Sigma-Aldrich) were applied at a 1:5,000 dilution for 1 h at room temperature. Immunoreactive proteins were visualised using Clarity Western ECL Substrate (BioRad) and a LAS-4000 imaging system (Fujifilm).

Antibodies used were: COX4 (1:500), α-Tubulin (1:10,000); Atg8 (1:1,000); and ref(2)P (1:1,000). Densitometric analysis was performed using Fiji software.

## 3. Results

### 3.1. Timing of CIV depletion differentially affects lifespan

To investigate the role of CIV on the adult lifespan of *Drosophila*, we employed RNAi against two different subunits of the mentioned respiratory complex: COX5B and COX4. We utilized the inducible tubulin-GeneSwitch (tubGS) driver to induce mitochondrial dysfunction starting at either the developmental stage or in adulthood (Figure 1A). We observed that initiating the knockdown of COX5B and COX4 during development resulted in massively short-lived flies (∼68% and ∼57% reduction for COX5B and COX4, respectively) (Figure 1B, C). However, when RNAi was activated only in adulthood, the reduction in lifespan was moderate for COX5B (∼20%) and even smaller for COX4 (∼11%) compared to controls with normal CIV levels. In both lines, flies where RNAi was induced only in adulthood were clearly long-lived compared to the short-lived lines where RNAi was induced during development. Furthermore, we observed a direct proportionality between the degree of lifespan reduction and the induction of mitochondrial dysfunction during development; the higher the concentration of RU-486 in the fly food, the more pronounced the shortening of adult lifespan was (Extended Figure 1A).

**Figure 1.**
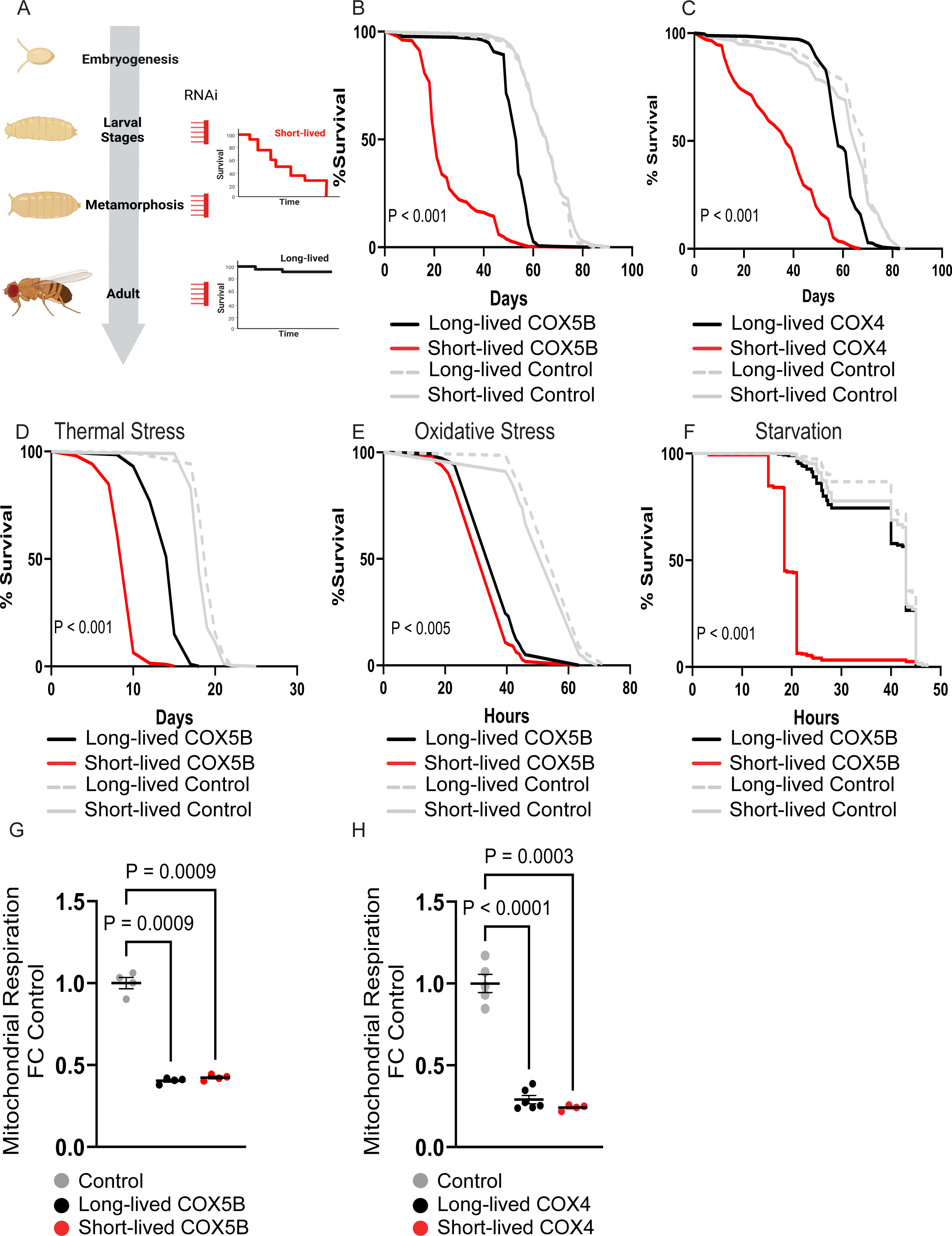
**Developmental depletion of Complex IV (CIV) drastically shortens adult lifespan.** (A) Schematic of the experimental design illustrating CIV depletion during both development and adulthood, or restricted to adulthood only. (B–C) Lifespan curves of the indicated genotypes. (D–F) Survival under the specified stress conditions. (G–H) Mitochondrial oxygen consumption rates in the indicated genotypes. Data are shown as mean ± SEM. Extended Figure 1. Supplementary analyses supporting Figure 1. (A) Lifespan curves of COX4 flies exposed to different RU-486 concentrations during development. (B–C) Mortality rates derived from Gompertz analysis for the indicated genotypes, with regression lines representing the fitted probability of death over time. (D–I) qPCR analysis of mRNA levels for selected genes across the specified genotypes and life stages. (J) Mitochondrial oxygen consumption rates in the indicated genotypes. Data are shown as mean ± SEM.

The mortality analysis of survival curves revealed opposing trends between short- and long-lived CIV-depleted flies. Short-lived models exhibited an earlier onset of mortality (increased α), but a markedly lower age-dependent mortality rate (reduced β) compared with both long-lived and control flies (Extended Data Figure 1B–C). In contrast, long-lived CIV-depleted flies showed accelerated age-related mortality (increased β) relative to both control and short-lived groups, despite an initial mortality rate comparable to controls in the case of COX5B depletion, and even delayed onset in the case of COX4 depletion. These findings indicate that adult-specific depletion of CIV accelerates ageing, in contrast to the effect observed upon CI depletion (Stefanatos, Robertson et al. 2025), supporting the notion that mitochondrial dysfunction promotes ageing when it simultaneously impairs ATP production and increases ROS generation. Notably, young long-lived CIV-depleted flies displayed no overt defects, including normal or delayed initial mortality, suggesting that detrimental effects emerge only later in life, when other cellular systems are already declining. As expected, RU-486 administration did not affect the survival of control flies with normal CIV levels (Fig. 1B–C).

Although not universally observed, changes in adult survival often correlate with resistance to external stresses (Wang, Kazemi-Esfarjani et al. 2004). To assess whether adult levels of CIV influence stress resistance, we subjected both short-lived and long-lived CIV-depleted flies to three distinct stress conditions: thermal stress (constant exposure to 32 °C), oxidative stress (administration of a 5% sucrose solution containing H₂O₂), and starvation stress (removal of food while maintaining access to water). Despite both groups harbouring mitochondria with aged-like features, their responses to stress differed markedly. Short-lived CIV-depleted flies exhibited consistently higher sensitivity than controls across all stress assays (Figure 1D–F). In contrast, the stress response of long-lived CIV-depleted flies was stress-type dependent: these flies survived starvation at levels comparable to controls, displayed intermediate sensitivity to thermal stress (more resistant than short-lived flies but less than controls), and showed a similar sensitivity to oxidative stress as the short-lived group (Figure 1D–F).

To verify that our experimental manipulations had the intended impact on CIV levels, we employed both quantitative PCR (qPCR) and high-resolution respirometry. As anticipated, we detected a decrease in mRNA levels of the target genes during development in experimental flies fed RU-486. Given that different RNAi constructs exhibit varied efficacies, it was not unexpected that a decrease in the target gene was observed in pupae for COX5B, while the decrease was already significant in L3 for the short-lived COX4 (Extended Figures 1D & E). Importantly, we observed no difference in the levels of either COX5B or COX4 in adults when comparing short-lived versus long-lived CIV-depleted flies (Extended Figure 1F &1G). Likewise, we detected no differences in mitochondrial oxygen consumption between short- and long-lived individuals (Figure 1G & 1H). Finally, we showed that the concentration of RU-486 used during development did not affect either COX5B mRNA levels or respiration per se in the control group (Extended Data Figure 1H–J).

### 3.2. Developmental depletion of CIV disrupts adult lipid metabolism, causing a continuous starvation-like state which saturates the autophagy recycling system

To gain insight into the impact of CIV depletion on lifespan, we conducted transcriptomics analysis of short- and long-lived flies in COX5B-depleted flies (Figure 2A). Principal Component Analysis (PCA) revealed a distinct separation between the short- and long-lived groups compared to the controls (Figure 2B). As expected, the two control groups with and without RU-486 feeding during development clustered together.

**Figure 2.**
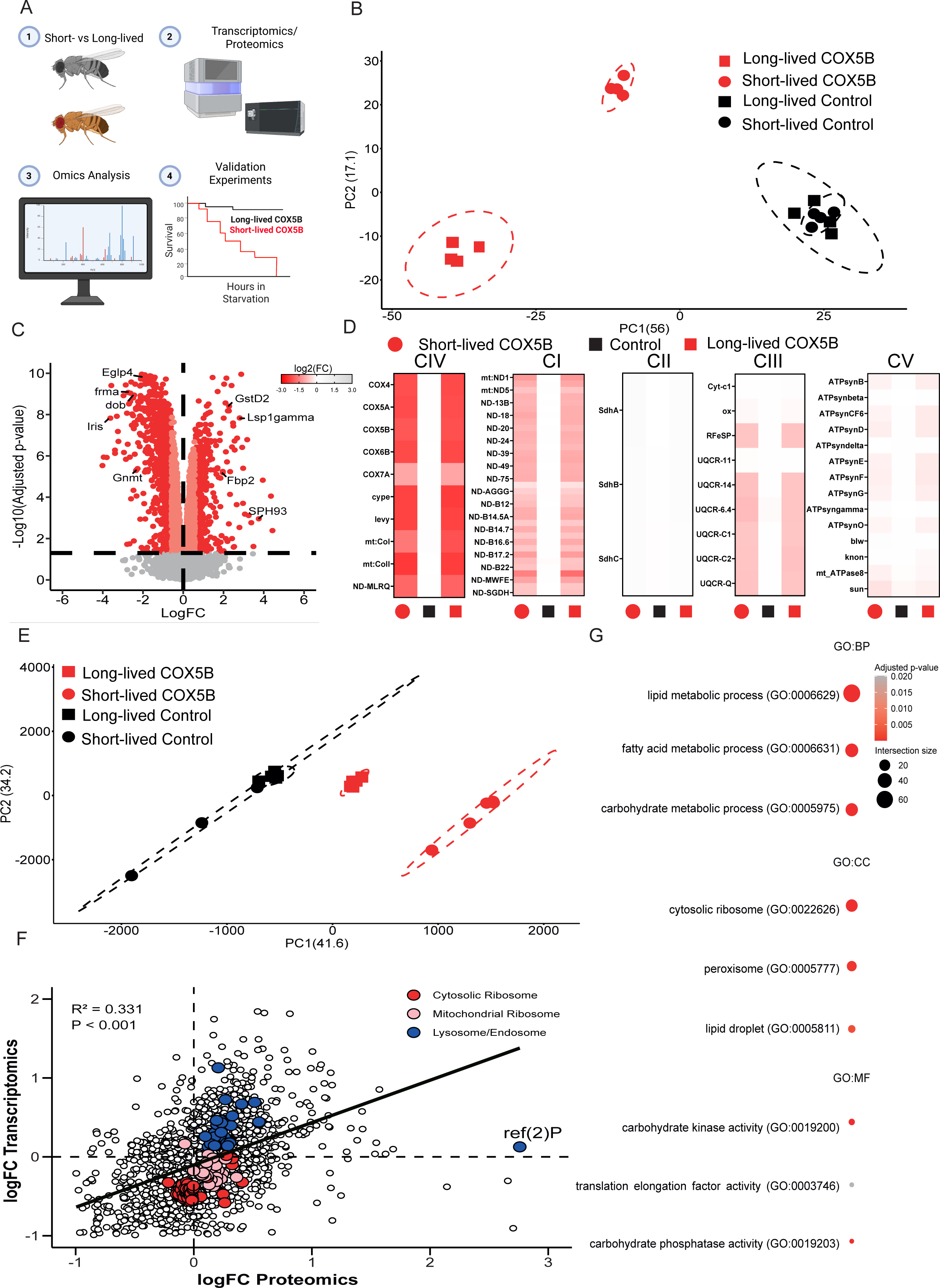
**Developmental CIV depletion induces adult misprogramming and a persistent state of metabolic starvation.** (A) Schematic of the omics analyses performed. (B) PCA plot of transcriptomic profiles showing clustering of short- and long-lived groups by genotype. (C) Volcano plot of differentially expressed genes (DEGs) between short- and long-lived COX5B flies. (D) Heatmaps showing expression levels of OXPHOS subunits in the indicated genotypes. (E) PCA plot of proteomic profiles showing clustering of short- and long-lived groups by genotype. (F) Scatter plot showing transcript and protein levels of the same genes and their correlation. (G) Dot plot of the most significantly enriched pathways identified by GO analysis. Extended Figure 2. Supplementary analyses supporting Figure 2. (A) Volcano plot of differentially expressed genes (DEGs) between short- and long-lived control flies. (B) Volcano plots showing the most significantly enriched genes in each indicated tissue based on FlyAtlas data. (C) Volcano plot of differentially expressed proteins (DEPs) between short- and long-lived control flies. (D) Representative confocal images of the fat body from the indicated groups: I, long-lived control; II, short-lived control; III, long-lived COX5B-depleted; IV, short-lived COX5B-depleted. Red indicates LipiTOX staining, and blue indicates DAPI. (E) Box-and-whisker plots showing triacylglyceride (TAG) levels across the indicated genotypes. Boxes indicate the median and interquartile range (IQR); whiskers denote minimum and maximum values. (F) Representative Western blot showing the indicated protein in the specified genotypes.

We identified 1,154 genes whose expression was significantly altered in short-lived COX5B-depleted flies compared with their long-lived counterparts, including 734 downregulated and 420 upregulated genes (Figure 2C). No genes were differently expressed when comparing control groups fed with RU-486 and those not fed with RU-486 (Extended Figure 2A), discarding that the differences observed are due to the presence of RU-486 during fly development. We utilised FlyAtlas 2 (Leader, Krause et al. 2018) to determine which fly tissues were most affected by CIV disruption during development (Extended Figure 2B). We observed that genes predominantly expressed in the carcass (primarily muscle), fat body, and heart were markedly downregulated in short-lived, COX5B-depleted flies (Extended Figure 2B). Conversely, genes enriched in the Malpighian tubules exhibited significant upregulation. A subset of genes primarily expressed in the larval fat body—typically absent or minimally expressed in adults—were among the most strongly upregulated in short-lived COX5B-depleted individuals. In contrast, genes characteristic of the adult fat body were, as noted, consistently among the most downregulated (Figure 2C). The resulting phenotype highlights tissue-specific disturbances in the muscle, heart and fat body of short-lived, CIV-depleted flies, resembling phenotypes previously described in short-lived CI-deficient individuals (Stefanatos, Robertson et al. 2025).

Next, we employed proteomics analysis to validate the depletion of the targeted CIV subunit. As anticipated, we observed a significant reduction in COX5B, along with all other CIV subunits, in both short- and long-lived flies, without significant differences between the two groups (Figure 2D). This further confirmed that observed differences between short- and long-lived flies are not caused by differences in the levels of adult CIV. Interestingly, depletion extended to many CI, CIII, and CV subunits, but not CII. This contrasts sharply with the effects observed following CI depletion, where only CI subunits are impacted (Stefanatos, Robertson et al. 2025). As it happened with the transcriptomic analysis, PCA separated short- and long-lived COX5B-depleted flies, but not the control groups (Figure 2E). Furthermore, only two proteins were identified as differentially expressed when comparing control groups, indicating that RU-486 minimally alters the fly proteome at the concentrations used in this study (Extended Figure 2C). Correlation analysis between transcriptomic and proteomic data revealed a weak, but statistically significant correlation when the expression levels of significantly altered genes and proteins in short-lived individuals were compared (Figure 2F). Accordingly, a significant number of genes and proteins exhibited concordant regulation, with 600 and 529 being commonly downregulated and upregulated, respectively. Additionally, a smaller yet significant number of proteins were upregulated despite reductions in their corresponding mRNA levels (295 proteins), while a minimal number displayed decreased protein levels despite upregulated transcripts (41 proteins).

Enrichment analyses of commonly downregulated genes and proteins revealed pathways crucial for energy production via carbohydrate and lipid metabolism, alongside diminished expression of proteins integral to peroxisomes and lipid droplets (Figure 2G). These alterations, consistently observed upon depletion of CI (Stefanatos, Robertson et al. 2025), suggest an induced starvation-like state attributable to the lack of proper differentiation of an adult fat body. To assess alterations in the former tissue, we examined its morphology and lipid content. LipidTox staining revealed a marked reduction in fluorescence exclusively in short-lived COX5B-depleted flies (Extended Data Fig. 2D), indicating impaired lipid accumulation. This suggests that, as observed for CI depletion (Stefanatos, Robertson et al. 2025), proper CIV levels are required for adult fat body differentiation. Consistently, triacylglycerol (TAG) quantification confirmed a significant reduction in lipid stores only in short-lived COX5B-depleted flies (Extended Figure 3E). Accordingly, these flies exhibited increased starvation sensitivity (Fig. 1F). None of these phenotypes were observed in control flies exposed to RU-486 during development (Figure 1F; Extended Figure 2D,E).

Notably, we observed a significant depletion in components of both cytoplasmic and mitochondrial ribosomes at the transcript level in short-lived COX5B-depleted flies (Figure 2F,G). Concurrently, a reduction in cytosolic ribosomal components and a moderate yet significant augmentation in mitochondrial ribosomal components were observed at the protein level (Figure 2F,G). While transcriptomic responses were conserved between CI- and CIV-short-lived phenotypes, proteomic alterations, particularly the divergent impacts on cytosolic versus mitochondrial translation, were characteristic of the CIV-short-lived flies (Stefanatos, Robertson et al. 2025) This indicates that similar phenotypic responses, such as energy deficits, are met with specific transcriptomic and proteomic adjustments depending on the OXPHOS complex affected. Finally, we observed an increase in the levels of proteins involved in quality control mechanisms, namely components of the lysosome and endosome (Figure 2F). In fact, one of the most upregulated proteins was ref(2)P, an autophagy receptor involved in the turnover of proteins and damaged organelles (Carroll, Otten et al. 2018). Its accumulation indicates impaired autophagic turnover, which was further confirmed by western blotting (Extended Figure 2F). The increase in ref(2)P was observed exclusively in short-lived COX5B-depleted flies, but not in their long-lived counterparts (Figure 2F and Extended Figure 2F) or in flies depleted of CI (Stefanatos, Robertson et al. 2025). These results indicate that alterations in CIV, but not CI, markedly affect autophagy; however, CIV depletion alone is insufficient to significantly alter the levels of either ref(2)P or Atg8b (Extended Figure 2F), as such changes only occur when depletion is initiated during development.

### 3.3 Targeted rescue during development improves energetic balance in adults, alleviates the blockade in autophagy and significantly extends adult lifespan

We tested whether mitochondrial functionality during development is required for an extended adult lifespan by complementing CIV activity exclusively in this period using the alternative oxidase (AOX) from *Ciona intestinalis* (Figure 3A). AOX has been shown to counteract the lethality caused by developmental CIV depletion (Kemppainen, Rinne et al. 2014); we therefore hypothesised that its expression would compensate for many of the alterations observed in short-lived COX5B adults. Accordingly, AOX expression restricted to development was sufficient to extend adult lifespan by ∼63% (Figure 3B) and starvation resistance by ∼90% (Figure 3C). This is remarkable given that AOX only partially restores electron flow and, when active, bypasses two proton-pumping sites (CIII and CIV) (Fernandez-Ayala, Sanz et al. 2009).

**Figure 3.**
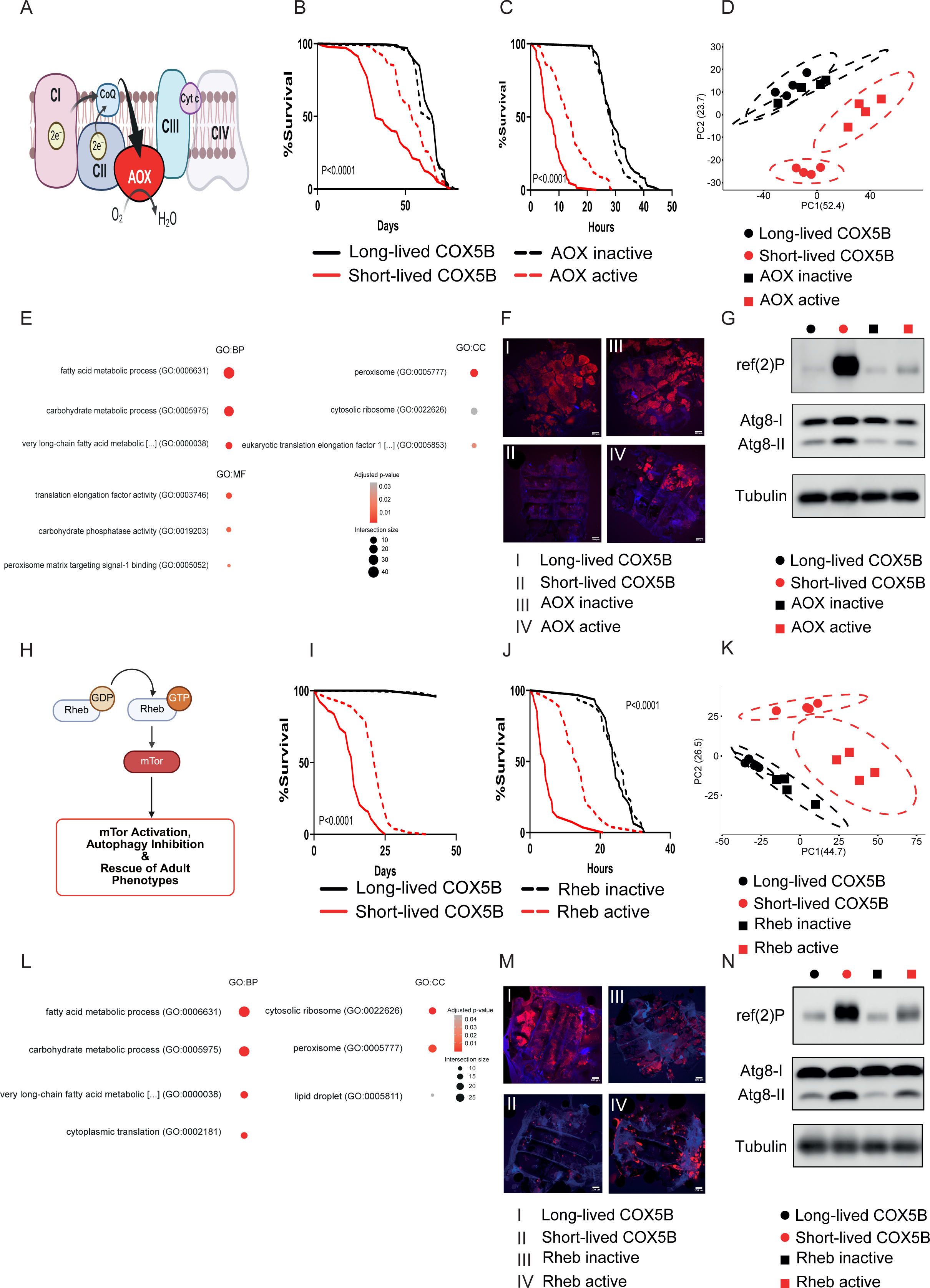
**Developmentally restricted expression of AOX or Rheb improves adult metabolic programming and extends lifespan.** (A) Schematic representation of how AOX partially restores mitochondrial respiration. (B) Lifespan curves of the indicated genotypes. (C) Survival under starvation. (D) PCA plot of transcriptomic profiles showing clustering of short- and long-lived groups by genotype. (E) Dot plot of the most significantly enriched pathways identified by GO analysis. (F) Representative confocal images of the fat body in the indicated groups: I, long-lived COX5B; II, short-lived COX5B; III, AOX inactive; IV, AOX active. Red indicates LipiTOX staining, and blue indicates DAPI. (G) Representative Western blot showing the indicated protein in the specified genotypes. (H) Schematic representation of how Rheb regulates mTor signalling and autophagy. (I) Lifespan curves of the indicated genotypes. (J) Survival under starvation of the indicated genotypes. (K) PCA plot of transcriptomic profiles showing clustering of short- and long-lived groups by genotype. (L) Dot plot of the most significantly enriched pathways identified by GO analysis. (M) Representative confocal images of the fat body in the indicated groups: I, long-lived COX5B; II, short-lived COX5B; III, Rheb inactive; IV, Rheb active. Red indicates LipiTOX staining, and blue indicates DAPI. (N) Representative Western blot showing the indicated proteins in the specified genotypes. Extended Figure 3. Supplementary analyses supporting Figure 3. (A) Heatmap showing transcriptional modules of genes downregulated or upregulated in short-lived COX5B knockdown flies, whose expression is restored or partially restored upon developmental AOX expression. (B) Box-and-whisker plots showing triacylglyceride (TAG) levels across the indicated genotypes. Boxes indicate the median and interquartile range (IQR); whiskers denote minimum and maximum values. (C) Heatmap showing transcriptional modules of genes downregulated or upregulated in short-lived COX5B knockdown flies, whose expression is restored or partially restored upon developmental Rheb expression. (D) Box-and-whisker plots showing TAG levels across the indicated genotypes, as in (B).

To investigate how developmental AOX expression extends lifespan in CIV-depleted flies, we generated transcriptomes from these flies and the corresponding controls. PCA revealed a clear separation between short-lived COX5B-depleted flies with or without AOX, whereas no distinction was observed in their long-lived counterparts (Figure 3D). Genes significantly rescued by developmental AOX expression (Extended Figure 3A) were enriched for fatty acid metabolic process (GO:0006631), carbohydrate metabolic process (GO:0005975), and cytosolic ribosome (GO:0022626) (Figure 3E). These transcriptional changes likely contribute to the lifespan extension and enhanced starvation resistance observed in AOX flies, although the rescue was incomplete, with many metabolic and cellular genes remaining unaffected, consistent with the partial nature of AOX complementation. In line with these results, AOX expression partially restored lipid and TAG levels in both the fat body (Figure 3F) and whole flies (Extended Figure 3B), which may underlie the improved starvation resistance. Finally, developmental AOX expression ameliorated one of the most prominent molecular defects of short-lived COX5B-depleted flies, impaired autophagy (Figure 3G), likely reflecting the increased fat reserves and alleviation of the chronic starvation state.

Having shown that partial complementation of mitochondrial function during development can rescue several adult phenotypes of short-lived COX5B flies, we next asked whether developmental interventions that do not restore CIV activity could also improve adult outcomes. We hypothesised that developmental mitochondrial dysfunction leads to a misprogramming of adult metabolism that might be corrected by targeting downstream pathways regulated by mitochondrial activity. To test this, we overexpressed Rheb specifically during development (Figure 3H). Rheb, an activator of Target of Rapamycin (TOR), suppresses autophagy when active (Hall, Grewal et al. 2007). Given that COX5B-depleted flies experience chronic starvation and defective autophagy, we reasoned that developmental Rheb activation might improve starvation resistance and extend adult survival.

In agreement with the above hypothesis, developmental Rheb expression extended adult survival and starvation resistance by ∼64% and ∼180%, respectively (Figure 3I and 3J). RNA-seq analysis revealed that transcriptomes of short-lived COX5B flies with or without Rheb expression separated by PCA, whereas controls clustered together (Figure 3K). Genes downregulated in short-lived COX5B flies but restored or partially restored by Rheb expression (Extended Figure 3C) were enriched for fatty acid metabolic process (GO:0006631), carbohydrate metabolic process (GO:0005975), and cytosolic ribosome (GO:0022626), closely resembling the pattern observed with AOX expression (Figure 3L). Like AOX, developmental Rheb expression increased lipid content in both the fat body and whole flies (Figure 3M and Extended Figure 3D) and restored autophagy markers towards the levels of long-lived COX5B controls (Figure 3N).

In conclusion, our experiments show that either partial complementation of mitochondrial function during development or targeting downstream pathways altered by mitochondrial dysfunction is sufficient to reprogramme metabolism, enhance lipid storage, alleviate chronic starvation, and extend adult survival in COX5B short-lived flies.

### 3.4 Levels of OXPHOS components are established during development and remain stable during the adult lifespan

Having shown that adult survival can be extended by either restoring mitochondrial function or compensating for its downstream consequences during development, we next tested whether re-establishing mitochondrial function in adulthood alone could rescue adult lifespan and stress resistance. To this end, we expressed an RNAi-resistant version of COX5B carrying silent mutations (smCOX5B; Figure 4A). Expression of smCOX5B in COX5B-depleted flies during adulthood failed to restore mitochondrial respiration (Figure 4B). In contrast, developmental expression of smCOX5B using the inducible GeneSwitch system enabled progression to the pupal stage under otherwise lethal conditions (Extended Figure 4A). However, no flies eclosed. While unexpected, this result was not entirely surprising, as the absence of feeding during the pupal stage limits the capacity of the rescue construct to support the assembly of a functional CIV.

**Figure 4.**
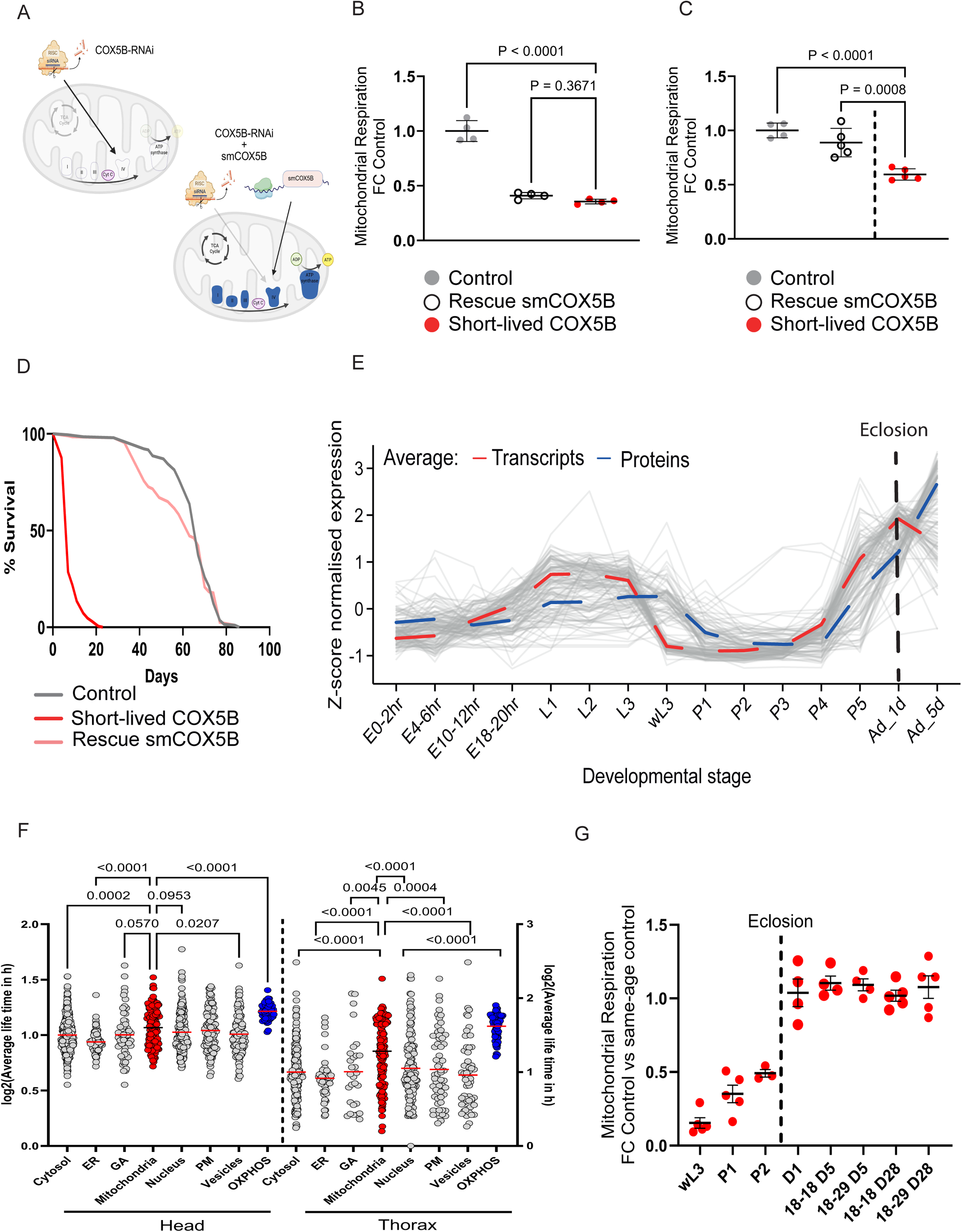
**OXPHOS levels are fixed during development and remain unaltered during adulthood.** (A) Schematic illustrating the strategy to restore CIV levels by expressing a copy of COX5B carrying silent mutations (smCOX5B). (B–C) Mitochondrial oxygen consumption rates in the indicated genotypes. (D) Lifespan curves of the indicated genotypes. (E) Temporal expression profiles of OXPHOS subunits at transcript (red) and protein (blue) levels throughout development and adulthood. Shaded lines indicate individual subunits; dashed lines represent the average. E=embryo, L=larvae, P=pupae, Ad=adult, hr =hour, d =day. The dash line indicate the time of eclosion. (F) Scatter plot showing the average lifetime in hour (log scale) of proteins localised to the indicated organelles or cellular compartments. Mitochondrial proteins are shown in red and OXPHOS proteins in blue. ER = Endoplasmic Reticulum, GA = Golgi Apparatus, PM = Plasma Membrane. (G) Mitochondrial oxygen consumption rates in flies with ND-18 RNAi induced at different developmental stages. The x-axis indicates the timing of RNAi induction: wandering L3 (wL3), pupae day 1 (P1), or pupae day 2 (P2). D1 flies were developed at 18°C and respiration was measured on day 1. Groups labelled as 18–18 D5, 18–29 D5, 18–18 D28, and 18–29 D28 correspond to flies in which RNAi was induced at D1 and respiration measured at day 5 or 28 of adulthood, raised continuously at 18 °C or shifted to 29 °C post-eclosion. Data are presented as mean ± SEM. Extended Figure 4. Supplementary analyses supporting Figure 4. (A) Representative images showing developmental progression in short-lived COX5B knockdown flies fed increasing doses of RU-486, with or without the smCOX5B rescue construct. (B) Representative images showing that COX5B knockdown under the strong daGAL4 driver prevents eclosion, which is rescued by expression of smCOX5B but not GFP. (C) Quantification of eclosion rates in COX5B knockdown flies with or without the smCOX5B construct, using da-GAL4 combined with tub-GAL80 to temporally control RNAi induction. (D) Representative Western blot showing protein levels in the indicated genotypes. (E) Scatter plots showing mitochondrial oxygen consumption rates in the indicated genotypes. (F) Scatter plots showing mg of protein per fly in the indicated genotypes. Data are presented as mean ± SEM.

To maintain rescue transgene expression during pupation, we employed a ubiquitous daughterless-GAL4 (daGAL4) driver. Notably, daGAL4 drives stronger RNAi expression than tubGS under moderate RU-486 concentrations, as used here. As expected, flies expressing only the RNAi construct did not eclose, whereas co-expression of smCOX5B enabled adult eclosion (Extended Figure 4B). To determine whether mitochondrial respiration was restored in adult flies, we repeated the experiment co-expressing daGAL4 alongside a tubulin-GAL80 temperature-sensitive repressor to moderate RNAi expression. Under these conditions, fewer than 15% of flies expressing COX5B RNAi successfully eclosed, whereas over 70% eclosed in the rescue control (Extended Figure 4C). To verify whether impaired eclosion was linked to reduced CIV levels, we assessed COX4 protein abundance in pupae by western blotting. We observed a marked depletion of COX4 in RNAi-expressing pupae, which was rescued by the presence of the control construct (Extended Figure 4D).

We performed high-resolution respirometry on flies co-expressing the RNAi and the smCOX5B rescue construct since development. No differences in mitochondrial respiration were detected between these flies and controls, either before or after normalising for total protein content (Figure 4C and Extended Figure 4E). Interestingly, in the minority of eclosing individuals (<15%) carrying only the RNAi construct, respiration was significantly reduced prior to normalisation but not after, indicating that the observed effect was due to a substantial reduction in total protein per fly (Extended Figure 4E-F). These RNAi-only flies also exhibited markedly shortened lifespans that was almost completely rescued by the presence of the rescue construct (Figure 4D). Together, these results confirm that the smCOX5B construct successfully rescues the developmental defects in mitochondrial respiration, eclosion, and adult lifespan caused by strong CIV depletion during development.

The former results suggested that CIV subunit levels are established during development and subsequently remain stable throughout adulthood. To investigate this hypothesis further, we analysed the expression profiles of mRNAs encoding OXPHOS components and their corresponding proteins during both developmental and adult stages using publicly available datasets from FlyBase. Our analyses revealed a distinct expression pattern characterised by two prominent peaks (Figure 4E). The first peak occurred at the third-instar larval stage (L3), followed by a notable decline in both transcript and protein levels during the first stages of pupation. The second peak arose immediately after eclosion. Notably, transcript and protein expression were closely correlated throughout developmental stages but began to diverge post-eclosion: protein levels remained elevated while transcript abundance progressively declined. This divergence, characterised by high protein stability alongside declining mRNA expression, aligns with trends previously reported to persist during ageing by multiple independent studies (Pletcher, Macdonald et al. 2002, Vincow, Thomas et al. 2021).

To test the hypothesis that OXPHOS protein levels are fixed during development and remain unchanged during adulthood, we first analysed protein half-lives in fly heads and thoraxes using two independent datasets (Vincow, Thomas et al. 2021, Ren, Xu et al. 2023). Supporting our hypothesis, we found that mitochondrial proteins exhibit significantly longer half-lives than proteins from other organelles, with OXPHOS subunits displaying the longest lifespans of all groups (Figure 4F). Finally, we performed a time-specific knockdown of a CI subunit (ND-18) at various developmental stages. We selected CI to avoid confounding effects, as CIV depletion can destabilise other respiratory complexes (Figure 2D), whereas CI depletion remains largely restricted to its own complex (Stefanatos, Robertson et al. 2025). We observed that CI knockdown prior to the third day of pupation at 18 °C led to a pronounced reduction in CI-linked respiration. In contrast, adult-specific depletion of CI had no measurable effect on respiration in either young or aged flies (Figure 4G). Notably, even after 28 consecutive days of ND-18 knockdown, respiration levels remained indistinguishable from those in control animals. This demonstrated that the same RNAi construct, which effectively depleted the target OXPHOS subunit and consequently reduced mitochondrial respiration when expressed during development, failed to affect OXPHOS when its expression was confined to adulthood.

## 4. Discussion

Our investigation sheds light on the central role of CIV in lifespan regulation, underscoring two distinct "windows of opportunity" where its influence is exerted. The initial window appears during developmental stages, with a secondary one manifesting during senescence. We demonstrate that the adult lifespan is reduced in proportion to the extent of CIV disruption experienced during development (Figure 1). The dramatic effect of CIV disruption during development contrasts with the moderate lifespan reduction observed when CIV depletion is limited to adulthood.

We previously reported that CI depletion during development produced a similar phenotype (Stefanatos, Robertson et al. 2023) . However, “adult-limited” CI depletion did not shorten survival. This finding suggests that different respiratory complexes play distinct roles in regulating adult lifespan, although their effects when targeted during development are similar. The observation that adult-restricted depletion of CIV, but not CI, shortens lifespan is consistent with the fact that CIV depletion induces aged-like mitochondrial features—reduced respiration and elevated ROS production—whereas CI depletion affects only mitochondrial respiration (Graham, Stefanatos et al. 2022). Accordingly, a recent paper (Lesner, Wang et al. 2022) reports that CIV is essential to maintain metabolic balance and protein homeostasis in the adult mouse liver, but CI is dispensable, which agrees with our observations.

The mortality analysis of lifespan reveals a distinct death rate pattern in short-lived flies. They began to die earlier, but their death rate was slower, suggesting a distinctive interplay between developmental mitochondrial function, adult survival, and death dynamics. Our new findings, along with previous ones regarding CI (Stefanatos, Robertson et al. 2025), highlight the significance of maintaining mitochondrial function during development, as it potentially sets the course for future health and longevity. This concurs with earlier research in worms and flies, where an increase in ROS levels during development extended lifespan (Obata, Fons et al. 2018, Bazopoulou, Knoefler et al. 2019). In our case, the primary role of ROS can be discounted, as we and others have observed contrasting effects of CI and CIV depletion on them (Graham, Stefanatos et al. 2022, Granat, Knorr et al. 2024). The role of CIV during adulthood aligns more closely with the expectation that carrying “aged mitochondria” throughout life is detrimental to longevity (Lopez-Otin, Blasco et al. 2023), as evidenced by a reduction in median lifespan and an accelerated mortality rate compared with controls who maintain normal CIV levels. However, it is noteworthy that young CIV-depleted flies, even when respiration was reduced more severely than in naturally aged flies (i.e. >50% (Scialo, Sriram et al. 2016)), showed no detectable defects during the reproductive period—the most evolutionarily important stage of the fly’s life—and began to die at the same time, or even later, than controls (Figure 1). This observation supports the notion that mitochondrial dysfunction alone is not the primary cause of ageing, but rather a consequence and a major contributor to its acceleration when other cellular systems fail in parallel. Experiments selectively depleting CIV in old individuals or restoring CIV specifically in old CIV-depleted flies, would help to clarify the precise role of mitochondria in this process. However, due to the high stability of OXPHOS complexes in adult flies, such experiments are currently not feasible with the available genetic tools.

Molecular characterisation of the short-lived COX5B knockdown flies (Figure 2) revealed that developmental depletion of CIV profoundly reprogrammed the adult transcriptome and proteome. This included widespread dysregulation of energy metabolism, with prominent alterations in lipid and carbohydrate homeostasis and cytosolic translation— phenotypes also observed in short-lived CI-deficient flies (Stefanatos, Robertson et al. 2025). These shared alterations in adult metabolism certainly contributed to the reduced lifespan observed in both CI- and CIV-depleted animals, reinforcing the notion that development is the critical life stage during which mitochondrial function exerts the greatest influence on adult lifespan in *Drosophila*.

Notably, the effects of developmental CIV depletion were most pronounced in the fat body. Given the fat body’s central role in energy storage and metabolic regulation, the reduced starvation resistance observed in short-lived CIV-depleted flies does not come as a surprise. As previously reported for short-lived CI-deficient animals (Stefanatos, Robertson et al. 2025), CIV-depleted flies displayed a markedly reduced capacity of the fat body to store lipids (Figure 3). Impaired fat body function, characterised by defective lipid storage and reduced trehalose mobilisation, likely contributes to a chronic starvation-like state. As a result, peripheral tissues increase autophagy to generate alternative energy sources. However, chronic activation of cellular recycling pathways does not come without consequences. Due to sustained mitochondrial dysfunction, lysosomal degradation systems can become saturated and lose efficiency (Fernandez-Mosquera, Yambire et al. 2019), potentially explaining the accumulation of ref(2)P observed in short-lived flies with impaired fat body function, but not in long-lived counterparts with intact fat bodies. This distinction is critical, as it demonstrates that a lack of CIV *per se* (in adults) is not sufficient to trigger an autophagy defect that compromises the cellular recycling machinery. These results argue against a direct, universal link between mitochondrial deficiency and defective autophagy—a topic currently under active debate in the field (Tai, Wang et al. 2017)—and reinforce the concept that the timing of the defect is more critical than its magnitude.

We confirmed the importance of developmental stages in determining adult lifespan by implementing two distinct rescue strategies restricted to this period. Despite their mechanistic differences, both ectopic expression of AOX and overexpression of *Rheb* significantly extended adult lifespan, enhanced starvation resistance, increased lipid content in the fat body, and reduced ref(2)P accumulation—indicating partial restoration of the cellular recycling system (Figure 3).

As expected, neither intervention fully restored all phenotypes. AOX was employed to demonstrate that even a partial bypass of CIV (Fernandez-Ayala, Sanz et al. 2009), and thus mitochondrial function, was sufficient to rescue most defects observed in short-lived CIV-depleted flies. Rheb was selected to test whether the detrimental phenotypes in short-lived CIV-depleted flies could be mitigated independently of mitochondrial function.

As an activator of Tor, Rheb alleviated autophagy overload and lysosomal stress, consistent with its known protective effects in models of excessive autophagy caused by Atg1 overexpression (Scott, Juhasz et al. 2007) or Vps16A mutation (Takats, Varga et al. 2015). The observation that Tor activation is protective in this context suggests that the relationship between mitochondrial dysfunction and Tor signalling may be context-dependent, as Tor inhibition is protective in both mouse and fly models of CI dysfunction (Johnson, Yanos et al. 2013, Wang, Mouser et al. 2016). Supporting this view, recent studies have shown that rapamycin extends lifespan in some genetic backgrounds but not in others (Ibrahim, Bahilo Martinez et al. 2024), and that it is not always effective in treating mitochondrial dysfunction (Barriocanal-Casado, Hidalgo-Gutierrez et al. 2019).

The most unexpected finding of our study arose when we attempted to rescue CIV activity exclusively during adulthood using a construct carrying silent mutations. Unexpectedly, adult-specific expression of the rescue construct failed to restore mitochondrial respiration, although it functioned as expected when expressed during development, i.e. rescued respiration and extended adult survival (Figure 4). These results suggest that OXPHOS complex levels are established during development and remain stable throughout adulthood.

Multiple lines of evidence support the above, i.e. that OXPHOS complex abundance is programmed during development and maintained throughout adult life (Figure 4). First, transcriptomic and proteomic analyses reveal coordinated developmental regulation of OXPHOS components, with mRNA levels declining with age while protein levels remain stable after eclosion—consistent with a model in which OXPHOS proteins are synthesised early and persist thereafter. Second, turnover rates of mitochondrial proteins, particularly OXPHOS subunits, are exceptionally low in adult flies, with average half-lives suggesting they are not replaced during the entire adult lifespan (Figure 4).

Third, while RNAi targeting CI and CIV subunits is effective at the transcript level during both development and adulthood, reductions in protein levels and mitochondrial respiration are only observed when knockdown is induced during development (Figure 4).

Based on the above findings, we must conclude that in long-lived CIV-depleted flies, knockdown is initiated prior to eclosion despite the absence of RU-486 during development. The leaky expression of the tubGS driver (Scialo, Sriram et al. 2016) is likely sufficient to induce RNAi during the pupal stage, disrupting the assembly of both CI (Stefanatos, Robertson et al. 2025) and CIV (this study). Although this late depletion does not shorten lifespan in CI-deficient flies and only mildly affects CIV-deficient flies, it results in protein-level reductions comparable to those observed with earlier larval knockdown. These CI- and CIV-depleted flies likely represent the closest approximation to adult-only knockdown currently achievable in *Drosophila*, with onset in late pupal stages, thereby avoiding the detrimental effects of earlier mitochondrial dysfunction. Furthermore, our results show that mitochondrial respiration cannot be restored in adult flies if depleted during development, in contrast to observations in mice, where adult CI depletion can be reversed (Corrà, Checchetto et al. 2023).

From an evolutionary standpoint, it is logical for short-lived organisms such as *Drosophila* to prioritise reproduction over somatic maintenance and therefore exhibit greater mitochondrial plasticity during development than in adulthood. After eclosion, flies have only a few days to reproduce and pass on their genes. Consequently, a strategy that sets OXPHOS levels during development—based on environmental cues—and maintains them throughout adulthood (without spending precious resources on repair or turnover of the OXPHOS system) may be advantageous, even if it limits lifespan. In contrast, this strategy may be less adaptive in long-lived species, which instead appear to retain greater OXPHOS plasticity in adulthood, allowing turnover of damaged components and restoration of mitochondrial function following developmental perturbations, as seen in CI-depleted mice (Corrà, Checchetto et al. 2023).

Several factors may underlie the reduced OXPHOS plasticity in adult flies, including diminished transcription or translation, impaired protein import, or enhanced degradation of OXPHOS components by mitochondrial quality control systems. These mechanisms will require further investigation in future comparative studies between short- and long-lived species.

In summary, our findings provide key insights into the role of mitochondria in ageing. First, they support the existence of one or more critical developmental windows during which mitochondrial integrity is indispensable for long-term health, as even subtle impairments at this stage can markedly compromise eclosion or adult survival. Second, unlike CI, adult CIV activity appears to influence lifespan by accelerating ageing, although not necessarily causing earlier death. Third, although our results cannot be directly extrapolated to humans, they suggest that mitochondrial diseases should be targeted either during early development (Hyslop, Blakeley et al. 2016) or—if they arise in adulthood—through tissue-specific reprogramming strategies rather than global restoration of mitochondrial function (Jain, Zazzeron et al. 2019). Ultimately, our findings underscore the importance of accounting for developmental effects when assessing mitochondrial function in adults. This work highlights the intricate interplay between developmental and adult mitochondrial function in shaping lifespan, constrained by evolutionary pressures that prioritise reproductive success.

## 5. Acknowledgements

This research was supported by two BBSRC grants (BB/R008167/2 & BB/W006774/1) and a Wellcome Senior Research Fellowship (212241/A/18/Z) to A.S. R.T.K. was supported by grants from JSPS (20K22912; 25K18725), AMED (JP24gm6710024), Nakajima Foundation, Uehara Memorial Foundation, Senri Life Science Foundation and Sumitomo Foundation I.F.G was supported by an MVLS DTP studentship. V.I.K. acknowledges support by the H2020 Twinning project RESETageing (GA 952266).

## 6. Notes on authors’ contributions

B.C-V.R.Z. performed lifespan experiments, qPCR, high-resolution respirometry, western-blot, confocal imaging, and stress experiments.

R.Z. performed lifespan experiments, qPCR, high-resolution respirometry and western-blot experiments.

I.F-G. performed confocal imaging experiments and generated and analysed transcriptomics data. S.A. performed qPCR, lifespan experiments and provided technical assistance.

Y.Y., M.M. and M.F. generated and analysed proteomics data.

T.K. and V.I.K. conducted western-blot experiments.

R.S. and M.C. contributed to the design, supervision and /or analysis of data.

A.S. designed and supervised the project, performed data analysis, created the figures, and wrote the initial draft of the manuscript.

All authors contributed to analysing and discussing the results and revised the manuscript.

