## Supplementary_information for "Developmental Programming of Mitochondrial Function Limits Lifespan in Short-Lived Animals"

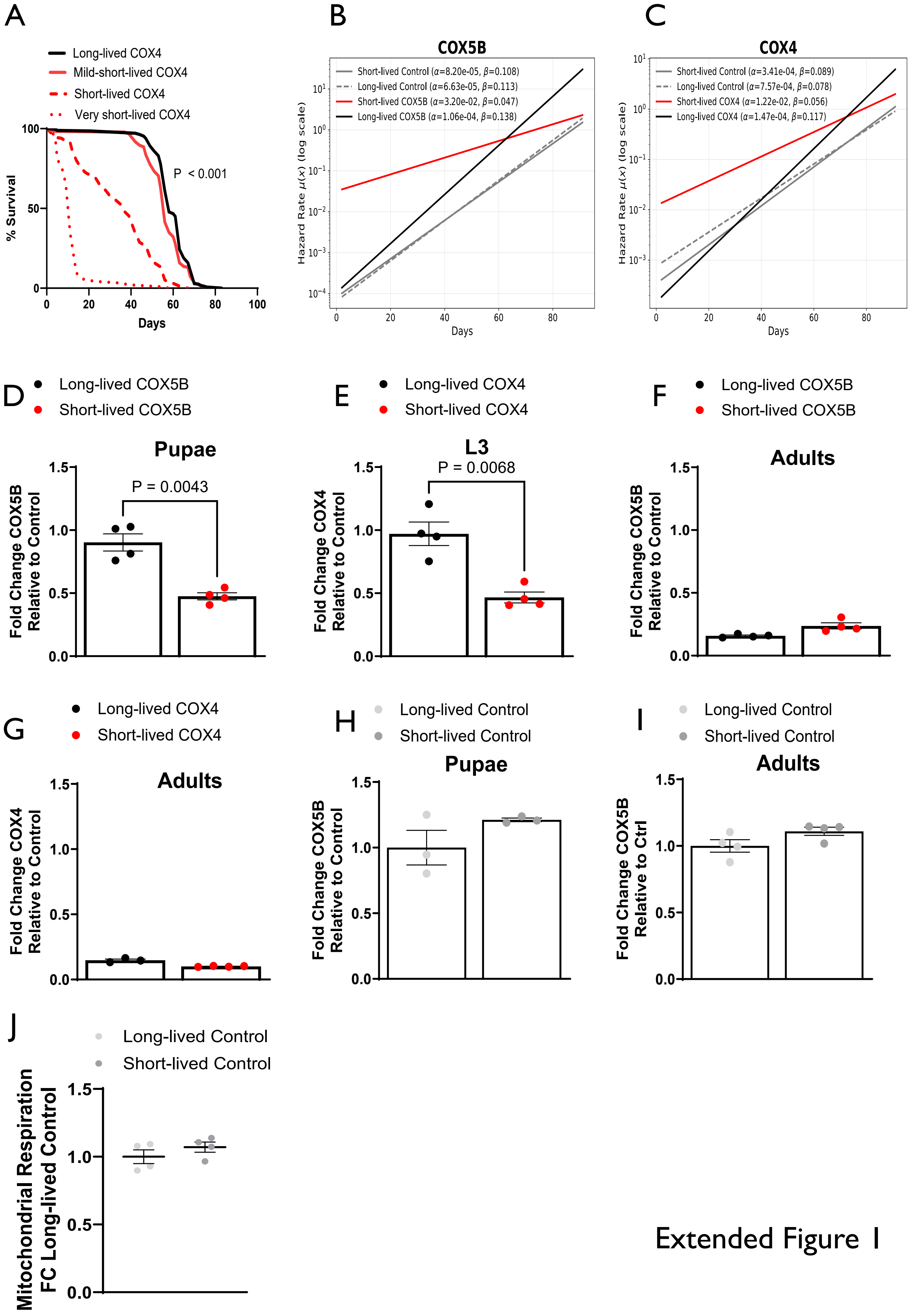


Extended Figure 1

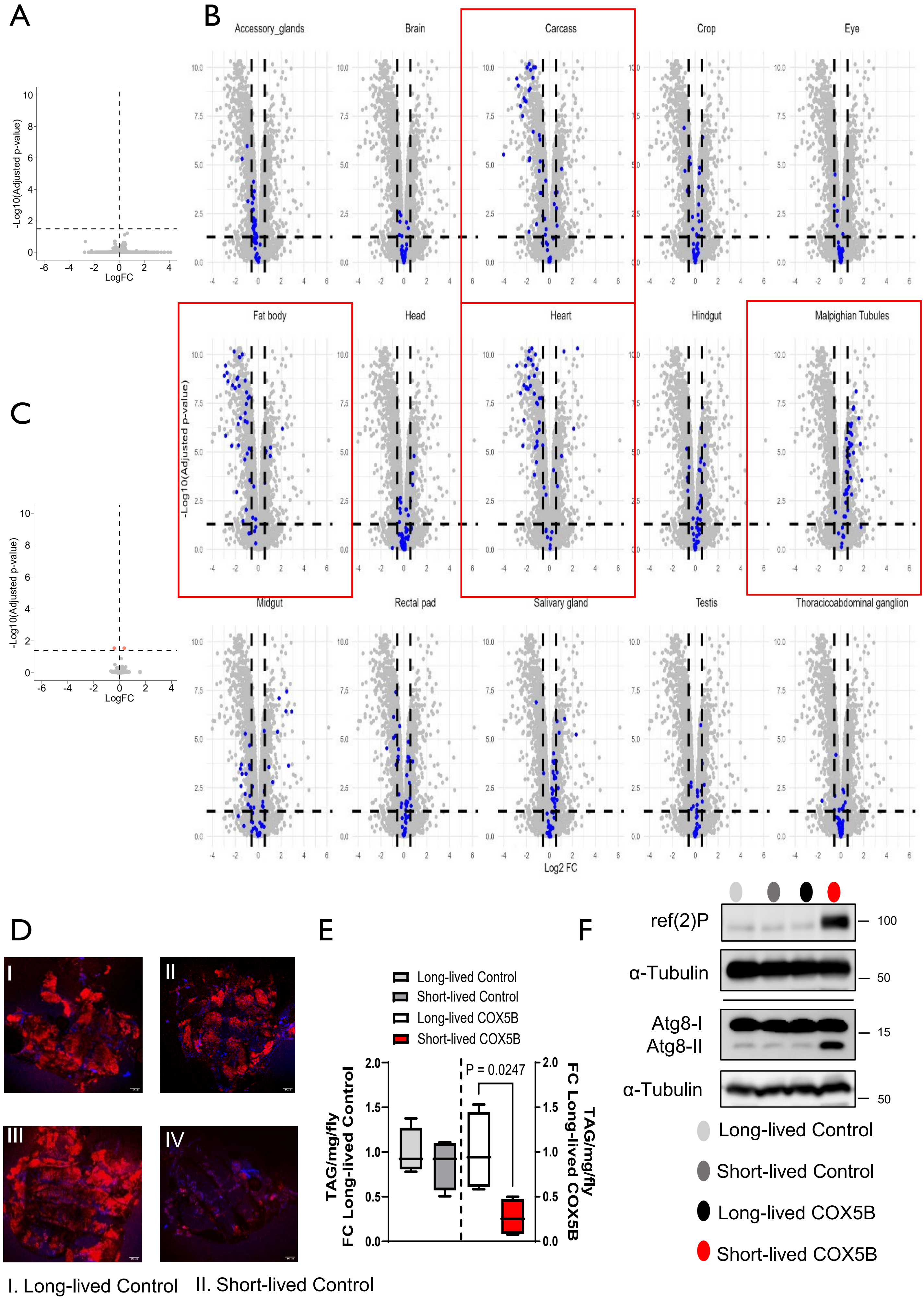


Extended Figure 2

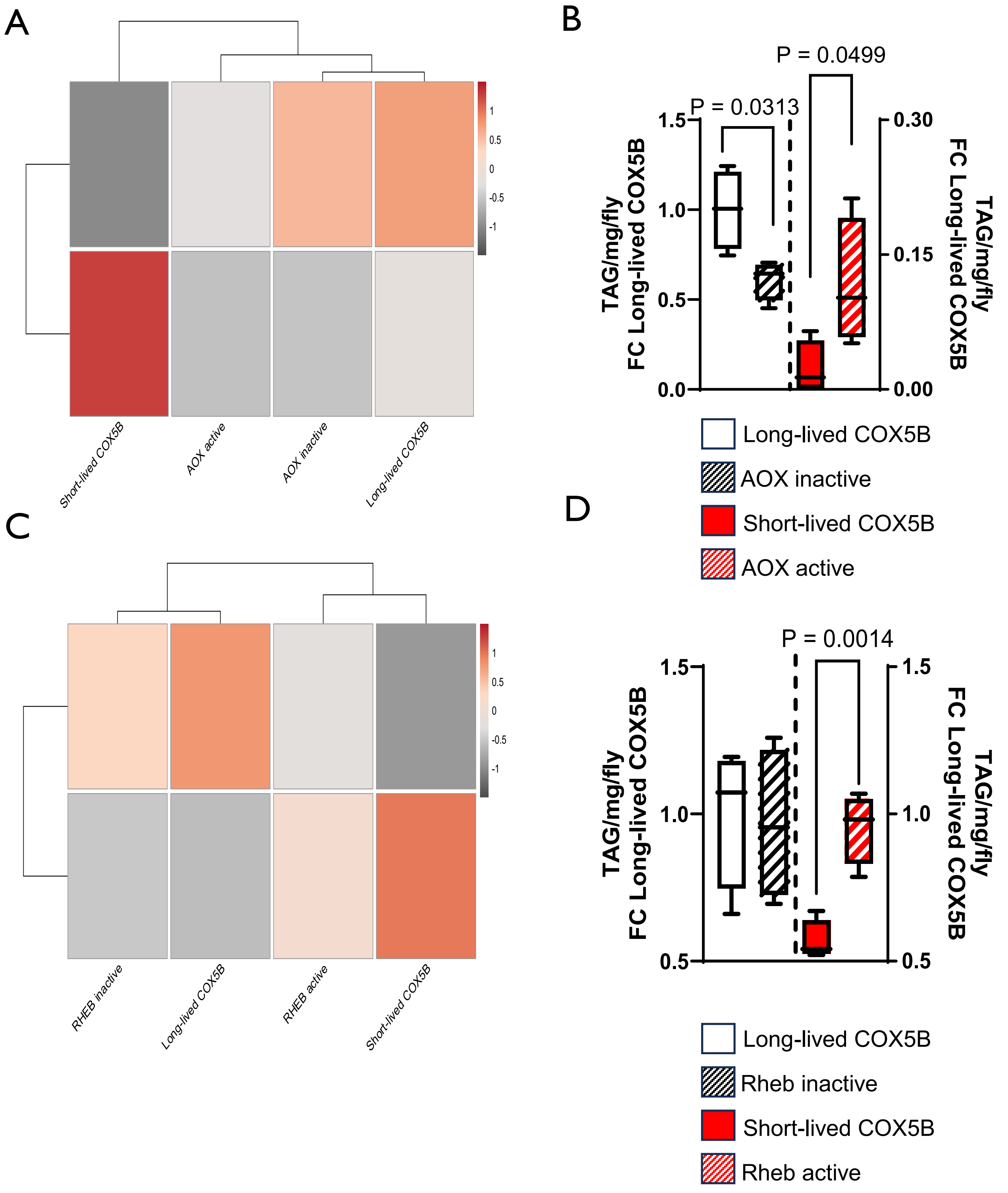


Extended Figure 3

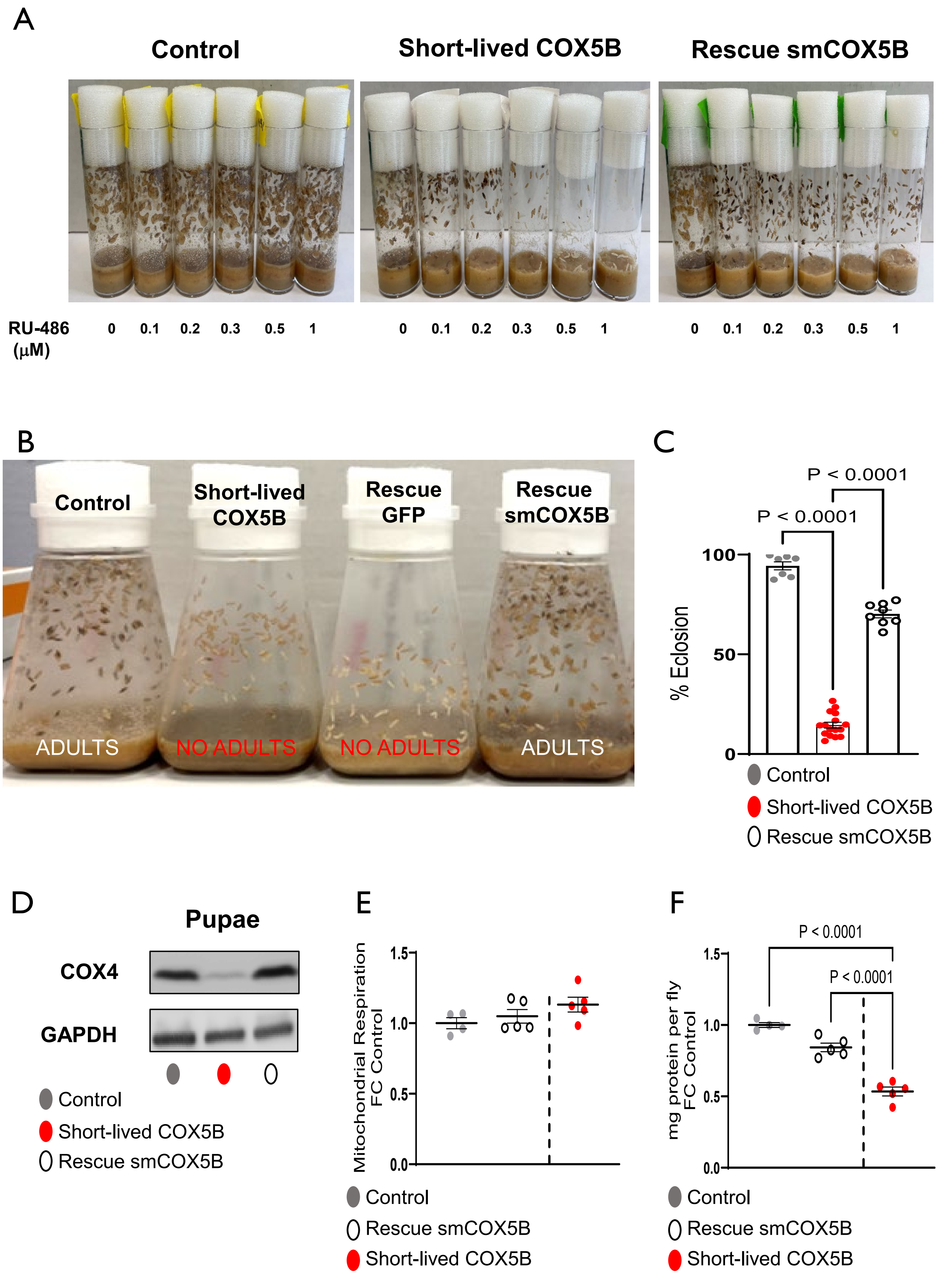


Extended Figure 4

### Supplementary Information

#### Developmental Programming of Mitochondrial Function Limits Lifespan in Short-Lived Animals

Beatriz Castejon-Vega[1], Ignacio Fernandez-Guerrero[1], Yizhou Yu[2,3], Rachel Zussman[1], Tetsushi Kataura[4], Rhoda Stefanatos[1,5,6], Sarah Allison[1], Davide D'Andrea[1], Mario Cordero[7], Milos Filipovic[1], Miguel Martins[3], Viktor I. Korolchuk[5], Alberto Sanz[1*]

[1] School of Molecular Biosciences, College of Medical, Veterinary and Life Sciences, University of Glasgow, G12 8QQ, Glasgow, United Kingdom.

[2] Healthspan Biotics Ltd, Milner Therapeutics Institute, CB2 0AW, Cambridge, United Kingdom.

[3] MRC Toxicology Unit, University of Cambridge, CB2 1QR, Cambridge, United Kingdom.

[4] Department of Neurology, Institute of Medicine, University of Tsukuba, Tsukuba, 305-8575, Ibaraki, Japan.

[5] Biosciences Institute, Faculty of Medical Sciences, Newcastle University, Campus for Ageing and Vitality, NE4 5PL, Newcastle upon Tyne, United Kingdom.

[6] Wellcome Centre for Mitochondrial Research, Faculty of Medical Sciences, Newcastle University, NE2 4HH, Newcastle upon Tyne, United Kingdom.

[7] Department of Molecular Biology and Biochemical Engineering, Pablo de Olavide University, 41013, Seville, Spain.

5

**Materials and Methods**

**Supplementary Table 1. Genotype and source information for fly lines used in this study**.

| Genotypes | Source |
|---|---|
| *w*[1118]*;; tubulin-GeneSwitch* | Gifted by Prof Scott Pletcher |
| *w*[1118]*;; daughterless-GAL4* | BDSC, BL#55851 |
| *w*[1118]*; tubulin-GAL80* | BDSC, BL#7019 |
| *w*[1118]*; UAS-COX5B* | VDRC, CG11015, v105769 |

| | |
|---|---|
| $w^{1118}$; *UAS-COX4* | VDRC, CG10664, v109338 |
| $w^{1118}$; *UAS-ND-18-IR* | VDRC, CG11015, v101489 |
| $w^{1118}$; *UAS-GFP* | The fly line was created for this study using BL24749. |
| $w^{1118}$;;*UAS-AOX;* | Described in (Fernandez-Ayala, Sanz et al. 2009) |
| $w^{1118}$;; *UAS-smCOX5B* | The fly line was created for this study using BL24749. |
| $w^{1118}$;; *UAS-Rheb* | The fly line was created for this study using BL24749. |
| $w^{1118}$; *UAS-Empty landing platform control* | VDRC, v60101 |
| $w^{1118}$; *UAS-Empty landing platform control* | VDRC, v60102 |
| $w^{1118}$; *UAS-Empty landing platform control* | A UAS-empty vector was inserted into the landing platform of BL24749. |

**Supplementary Table 2. Primer Sequences**

| Primer | Sequence |
|---|---|
| *Act88f* F | agggtgtgatggtgggtatg |
| *Act88f* R | cttctccatgtcgtcccagt |
| *AOX* F | attttctttggcttacttaatctcac |
| *AOX* R | caatttctggcgtctttca |
| *COX4* F | gagcccaccaacgagatcaa |
| *COX4* R | ttccagtctccctgctcctt |
| *COX5B* F | gctgcatctgcgaagaggat |
| *COX5B* R | caccagcttgaaccaatggc |
| *GAPDH* F | gacgaaatcaaggctaaggtcg |
| *GAPDH* R | aatgggtgtcgctgaagaagtc |
| *Rpl32* F | aggcccaagatcgtgaagaa |
| *Rpl32* R | tgtgcaccaggaacttcttgaa |
| *Rheb F* | tagccgtggtaatcctgct |

| | |
|---|---|
| Rheb | ctaccgatcagtgggcaaat |
